# Disinfection chemicals mode of action on the bacterial spore structure and their Raman spectra

**DOI:** 10.1101/2020.08.24.264440

**Authors:** Dmitry Malyshev, Tobias Dahlberg, Krister Wiklund, Per Ola Andersson, Sara Henriksson, Magnus Andersson

## Abstract

Contamination of toxic spore-forming bacteria is problematic since spores can survive a plethora of disinfection chemicals. It is also problematic to rapidly detect if the disinfection chemical was active, leaving spores dead. Robust decontamination strategies, as well as reliable detection methods to identify dead from viable spores, are thus critical. Vibrational detection methods such as Raman spectroscopy has been suggested for rapid diagnostics and differentiation of live and dead spores. We investigate in this work, using laser tweezers Raman spectroscopy, the changes in Raman spectra of *Bacillus thuringiensis* spores treated with sporicidal agents such as chlorine dioxide, peracetic acid, and sodium hypochlorite. We also imaged treated spores using SEM and TEM to verify if any changes to the spore structure can be correlated to the Raman spectra. We found that chlorine dioxide did not change the Raman spectrum or the spore structure; peracetic acid shows a time-dependent decrease in the characteristic DNA/DPA peaks and ∼20 % of the spores were degraded and collapsed; spores treated with sodium hypochlorite show an abrupt drop in DNA and DPA peaks within 20 minutes all though the spore structure was overall intact, however, the exosporium layer was reduced. Structural changes appeared over several minutes, compared to the inactivation time of the spores, which is less than a minute. We conclude that vibrational spectroscopy provides powerful means to detect changes in spores but it might be problematic to identify if spores are live or dead after a decontamination procedure.

## Introduction

A spore is an inactive seed-like form that some bacteria species can take to survive in a hostile environment. When faced with unfavorable conditions such as a lack of food, these bacteria will form spores to protect themselves in a process called sporulation.^1^ During sporulation, the vegetative cell copies its DNA and packs it within multiple protective layers, before finally bursting and releasing the completed spore into the environment.^2^ Spores are metabolically inactive (they do not grow or reproduce), but they contain the complete genome of the species, as well as the cellular machinery and receptors needed to germinate back into vegetative cells again upon contact with favorable conditions. As long as the bacteria remain in spore form, they can survive circumstances that would kill a vegetative cell. For example, spores can survive temperatures below freezing and above 100 ° C, exposure to strong acids (including stomach acid), antibiotics, ethanol, quaternary ammonium, and peroxide-based agents.^3^ Further, spores can survive in the environment for a very long time, easily into decades, such as with *B. anthracis* spores, unless decontaminated with strong chemical agents like formaldehyde of sodium hypochlorite.^4^

This extreme durability poses many problems for society as spores cause diseases in both humans and animals. For example, spores such as *B. cereus* and *C. perfringens* are common causes of food poisoning^5^ and *C. difficile* a cause of colitis diarrhea. Canned food can become contaminated with *C. botulinum* spores producing dangerous botulin toxin. In cases of infection by these bacteria, their durability puts extra strain on society due to the harsh decontamination methods needed to deal with them. ^6^ For example, hospital fabrics from *C. difficile* patients in hospitals cannot be washed with other fabrics as the spores will survive the high-temperature washes and contaminate all the fabric in the batch.^7^ Further, spores from the *Bacillus* genus such as those of *B. anthracis* present a potential biological warfare hazard since these spores are lethal and difficult to decontaminate.^8^

Several effective decontamination methods using chemicals exist. Chlorination is a popular approach; however, many strains of the *Bacillus* genus exhibit apparent resistances towards chlorination disinfection.^9^ Other proved effective decontamination chemicals for *Bacillus* strains are chlorine dioxide, sodium hypochlorite and peracetic acid.^10^ Even though these are indeed effective care must be taken since these compounds are unstable in regular conditions: chlorine dioxide and sodium hypochlorite decay and release chlorine (in itself a toxic gas), especially in sunlight, while peracetic acid decays back to acetic acid and hydrogen peroxide (which in turn decays to water and oxygen).^11^ While these chemicals decay into non-lethal components, they are initially very toxic to a plethora of organisms as well as human skin cells.^12^ Thus, to assess if a decontamination procedure was successful without overusing the sporicidal chemical, it is important to detect if a spore is dead or alive.

To identify dead from viable spores use of vibrational spectroscopy techniques such as Raman spectroscopy has been suggested.^13–15^ Using Raman it is possible to identify key molecular components of the spore. However, whether Raman methods can reliably differentiate between intact, damaged and inactivated spores has not been investigated thoroughly. Therefore, we use Laser Tweezers Raman spectroscopy (LTRS) and electron microscopy (SEM/TEM) imaging to determine whether spore inactivation affects the Raman spectrum and the spore structure. LTRS allows us to isolate and move a single spore (“trap it”) and simultaneously measure its Raman spectrum to gain insight into its molecular changes during chemical exposure; and EM imaging allows us to observe exterior and interior cell structure changes.

## Results and discussion

### Mapping vibration peaks of the spores using Laser Tweezers Raman spectroscopy

To measure the impact of the sporicidal chemicals on the spores’ surface structure as well their Raman peaks; we first assessed the vibrational peaks in the absences of chemicals on purified *B. thuringiensis* spores using LTRS. Our instrument and the measurement procedure are described in detail in the Experimental section. One of the main constituents and the most common biomarker for these spores is CaDPA, which accounts for approximately 20 % of the spore core weight.^16^ CaDPA is a protective component located in the core and essential for a spore’s resistance to heat. Figure 1A top panel shows the average Raman spectrum of (A) three dormant spores sequentially trapped in the LTRS, a spectrum of pure DPA (B) and DNA (C), and the spectra of purified sporicidal chemicals used in this study (D, E, F). In previous studies, it has been reported that there are no significant changes in the Amide bands >1400 *cm*^−1^ in the presence of chemicals. ^17^ We investigated the 1580 *cm*^−1^ Amide band for changes in Raman intensity (Figure S1), and saw no change for chlorine dioxide or peracetic acid. For sodium hypochlorite, the changes in Amide I band intensity followed the changes in DPA peak intensity. Therefore, to allow for fast acquisition rate of Raman signals we limited the spectral measurement range of our system to 600 - 1400 *cm*^−1^, which is where CaDPA and DNA peaks are to be found.^18^

**Figure 1:**
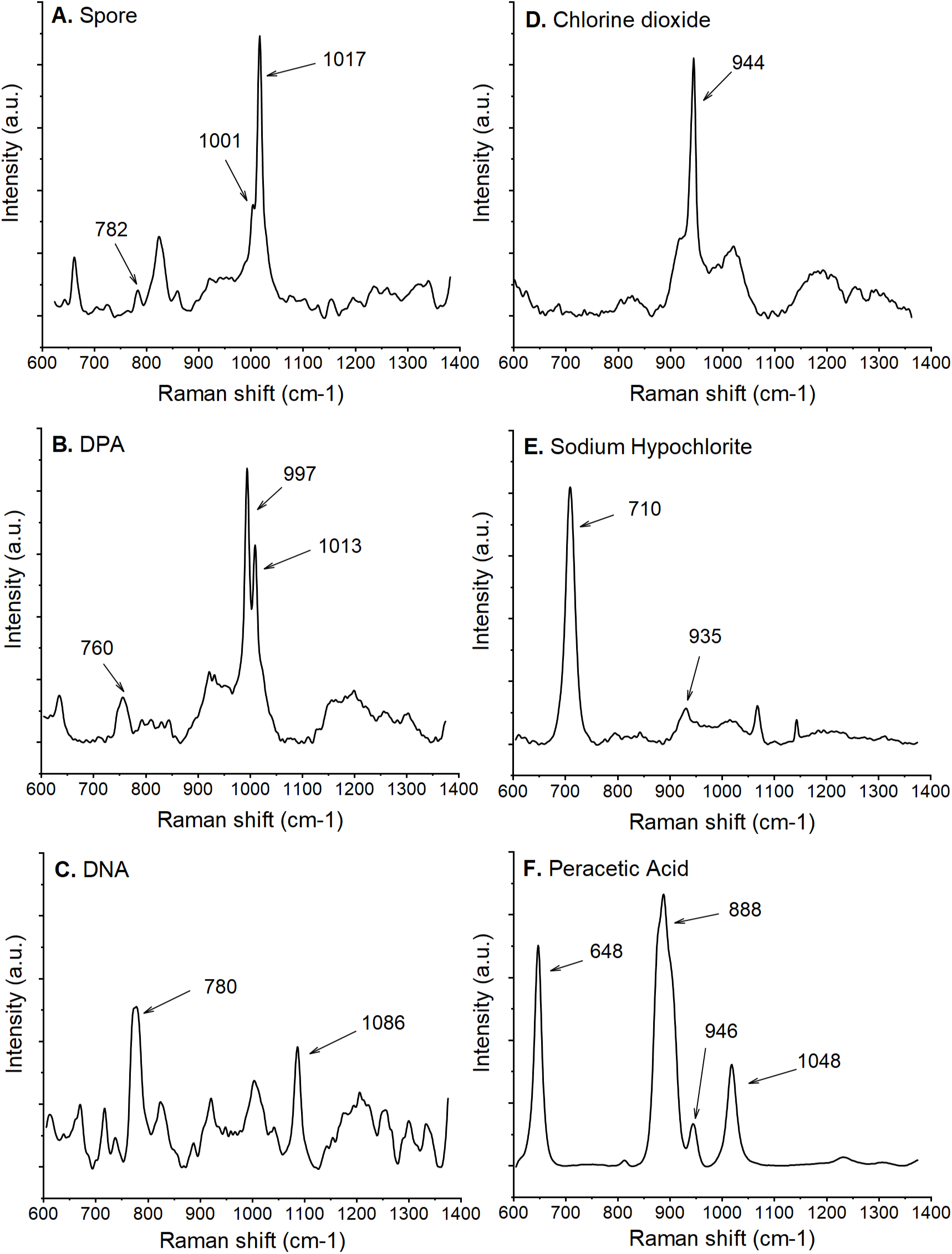
Raman spectra of spores, their major components and the sporicidal chemicals used. Each panel is an average of three spectra. A. is a typical spore Raman spectrum with the major peaks marked. The Raman spectra of spore components are included: DPA (B) and DNA (C). The Raman spectra of sporicidal chemicals are: chlorine dioxide (D), sodium hypochlorite (E), and peracetic acid (F) are shown.

Raman spectra of purified DPA and DNA is seen in Figure 1B and Figure 1C respectively. We marked the major CaDPA and DNA peak at 1017 *cm*^−1^ and 782 *cm*^−1^ in the top panel. These peaks are slightly shifted in the purified DPA and DNA spectra as in the purified solutions, since the bond length may be slightly different than when located in the spore. In particular, this is true with regards to pH and interactions with other spore components. For DNA, the 782 *cm*^−1^ peak is related to the O-P-O backbone or the cytosine ring breathing mode. Another peak that is clearly visible in the pure DNA Raman spectrum is at 1086 *cm*^−1^, related to the phosphodiester stretching peak. This peak, however, is difficult to observe in the whole spore spectrum.^19–21^ We also marked the phenylalanine peak, a major structural component of the spore, with the Raman peak at 1001 *cm*^−1^. Overall, the peaks observed using our LTRS is consistent with what is found in the literature using similar approaches.

### Chlorine Dioxide (DK-DOX) treatment does not affect DNA peaks whereas DPA peaks are reduced

Chlorine dioxide is a sporicidal chemical and it has been proven effective during decontamination of spores, without being very harmful to humans. However, the reported mechanism of action is not consistent in the literature, especially for DNA. For example, Zhu et al., reported that chlorine dioxide at concentrations higher than 100 ppm could damage DNA. At the same time, other publications suggest that the action of chlorine dioxide does not affect DNA directly.^22,23^ The concentrations used in these studies vary significantly from only a few ppm to several hundred ppm. To investigate the impact of chlorine dioxide on the Raman peaks and spore structure, we treated spores with chlorine dioxide at a concentration of 750 ppm (the upper end of the concentrations used in literature) and carefully recorded the time of exposure. Spores were trapped using our LTRS and the Raman spectrum of individual spores acquired. We found that the Raman spectrum of spores was not affected by incubation with chlorine dioxide, even though the concentration of 750 ppm (Figure 2A, is 75 times higher than the 10 ppm reported lethal to spores.^22^ Indeed, the spores were confirmed inactivated using a viability study (0 CFU on growth media plates, a 5-6 log reduction compared to control). We do note that there is a gradual minor decrease of signal intensity, 4 % for DPA and 6 % for DNA over the 30 minute measurement time. Due to the smaller absolute intensity of the DNA peak (Figure 1A), the changes in normalised intensity had more noise than in the DPA peak. Since these drifts observed are small and occurs over a 30 minute timescale, they do not influence the outcome of the results.

**Figure 2:**
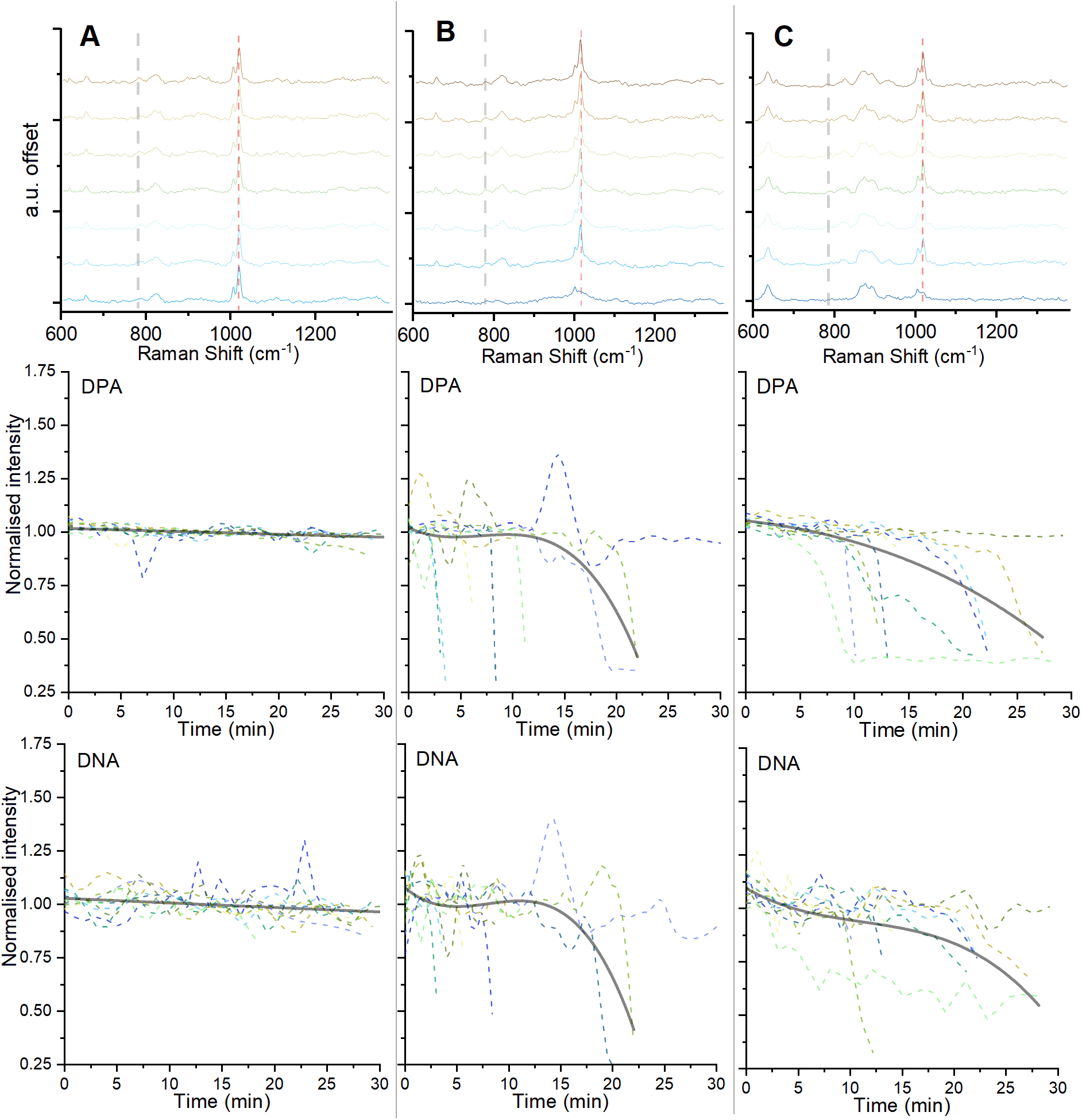
Graphs in the top row show change in the Raman spectrum of a single spore overtime where A) is with 0.075 % chlorine dioxide B) with 0.5 % sodium hypochlorite and C) with 1 % peracetic acid. The top data graph is the starting Raman spectrum (t=0), and each subsequent graph is a 4-minute increment. Dashed vertical lines indicate the major DPA (right) and DNA (left) peak, respectively. Middle and bottom rows show smoothed normalized intensities of the DPA and DNA peaks from ten individual spores vs. time. The solid grey lines are averages of all data.

Chlorine dioxide has been reported to react with and damage proteins, with cysteine and methionine in particular.^24–26^ However, we did not see a decrease in Raman peaks associated with proteins, such as the Amide I band. To see surface and internal structure differences between untreated and treated spores, we used SEM and TEM imaging. Examples of untreated spores in SEM and TEM are shown in 3A and 4A. We also added some additional TEM micrographs in Figure S2. Both SEM (n=20) and TEM (n=10) images of spores treated with chlorine dioxide (Figure 3B, Figure 4B) do not show any visible damage to the exosporium, spore coat or internal structure. Note that the specific Raman peak of chlorine dioxide at 944 *cm*^−1^ in Figure 1 is consistent with previously published data. ^27^ Although the peak rapidly decreased in intensity during the measurement, it disappeared in approximately 15 minutes indicating that chlorine dioxide disappeared from the solution, either by chemical decomposition into chlorine or by diffusing out. Taken all together, we conclude that spores inactivated by treatment with chlorine dioxide did not show any major changes in neither their Raman spectra as measure by LTRS, or exosporium and spore coat, as visualize using SEM, or internally as visualized using TEM.

**Figure 3:**
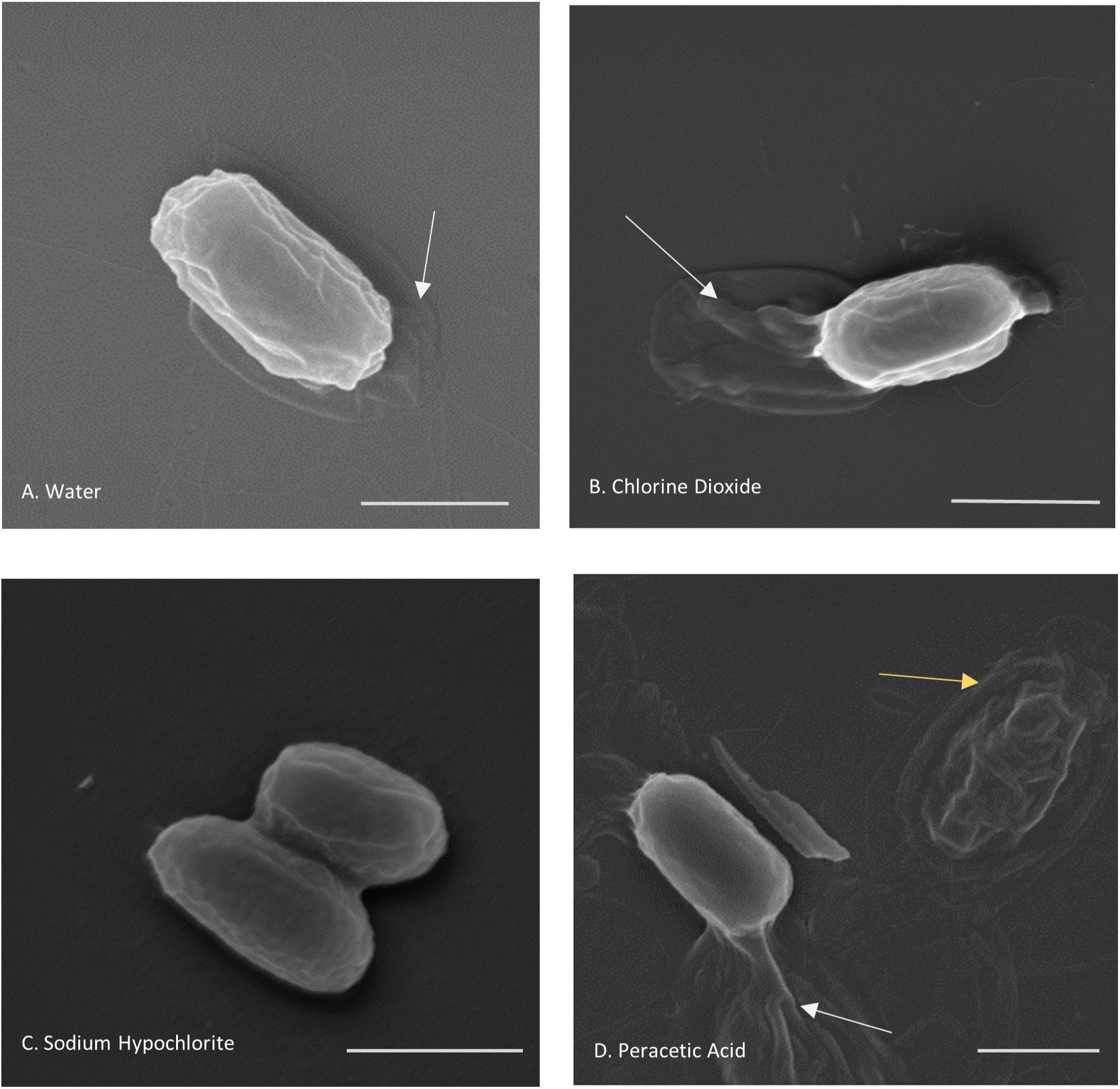
SEM of *B. thuringiensis* spores treated with different sporicidal chemicals, imaged at 50,000 magnification. The loose exosporium surrounding the spore (white arrows) can be seen for spores in A) purified and deionized water, B) chlorine dioxide and D) peracetic acid. No exosporium in spores treated with C) sodium hypochlorite is seen. A partially degraded spore is also seen in the image of D) peracetic acid treated spore (yellow arrow). Scale bars are 1 *μ*m.

**Figure 4:**
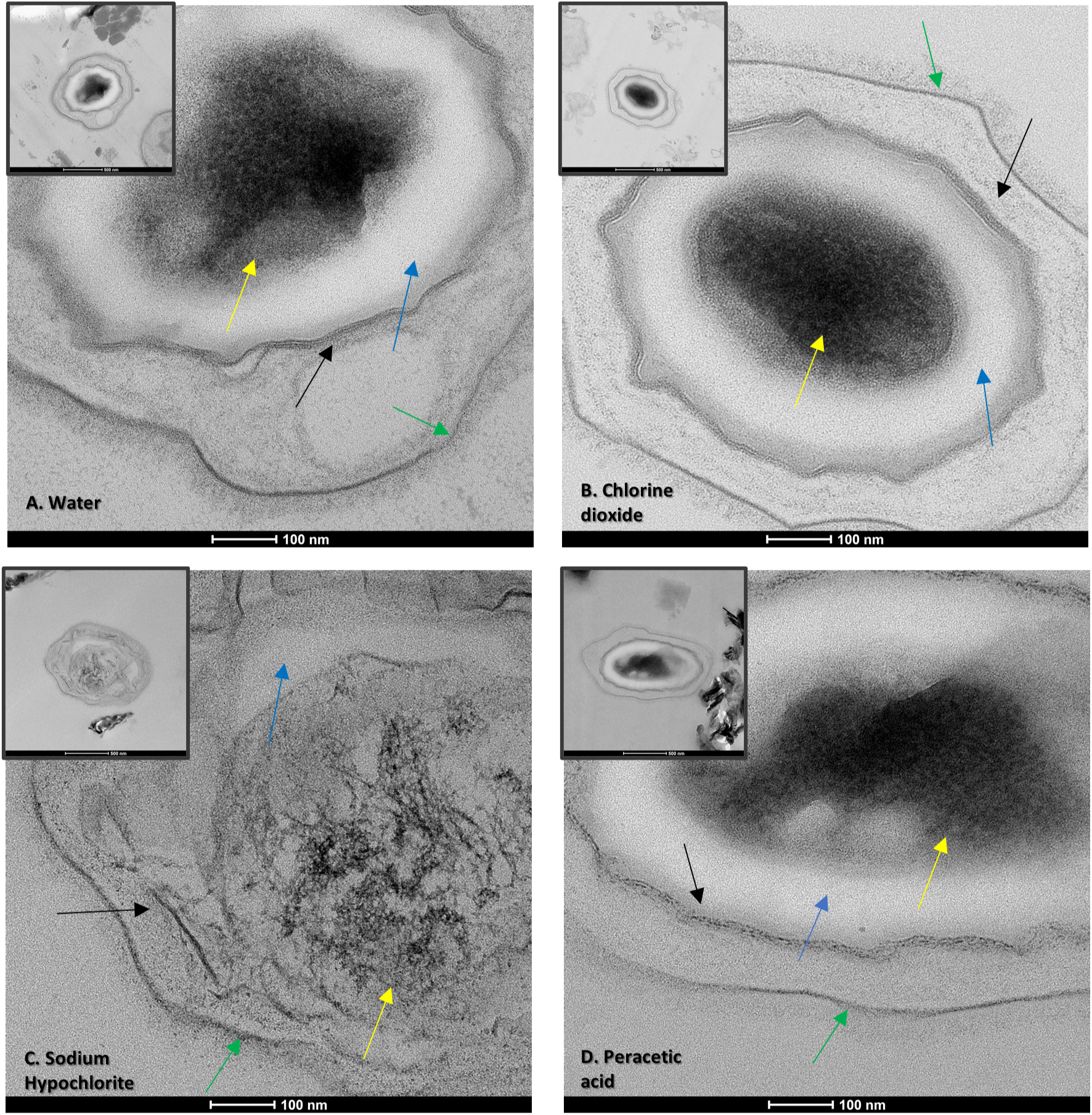
TEM of *B. thuringiensis* spores treated with different sporicidal chemicals, imaged at 73,000 magnification and 27,000 magnification (inset). We labeled the structural layers of the spore as follows. Green arrows: exosporium; Black arrows: spore coat; Blue Arrows: cortex; Yellow arrows: core. A) untreated spore in water B) Spores treated with chlorine dioxide appear no different to control. C) Spores treated with sodium hypochlorite are degraded, and the core is no longer electron-dense, indicating DPA release. D) Spores treated with peracetic acid have a fragmented spore coat.

### Effect of sodium hypochlorite

Next, we analyzed spores treated with a 0.5 % solution sodium hypochlorite (bleach) with a pH of 11.55. This concentration was previously reported to be sporicidal.^28^ Sodium hypochlorite is a cheap and prevalent decontamination agent that works by degrading organic material in several reactions: saponification of fatty acids and neutralisation and chloramination of amino acids.^29^ As such, hypochlorite causes oxidative damage to lipids, protein and DNA. We found that sodium hypochlorite causes the largest change in a spore’s Raman spectrum and strongly affects the DPA peak (Figure 2B), though there is a lag time before the decrease in the Raman associated DPA peaks begin. Though initially unaffected, the spores completely lose their major DPA peak at 1017 *cm*^−1^. The duration of the lag time before DPA loss varies, with the smallest being 4 minutes, and the largest being 22 minutes. One spore did not lose the DPA peak over the measurement time (Figure 2B). The speed at which a spore loses its DPA is rapid; in general the DPA signal can go from a full peak to a complete loss in less than a minute (2 minutes for the slowest spore). Finally, the Raman peak of DNA correlated with the DPA peak in both the time lag and rate of decrease.

When observed using SEM, the spores treated with sodium hypochlorite appear to have significant changes to the exosporium layer (Figure 3C), whereas the spore coat is still intact. Only 4 of the 22 spores imaged had a poorly visible exosporium, the remaining spores did not have it at all. The degradation of the exosporium is consistent with the mechanism of action of hypochlorite, with the most damaged structure being the outer layer.^29^ The loss of DPA in hypochlorite treated spores is interesting. It was previously reported that spores treated with hypochlorite do not lose their DPA store from treatment, but they do germinate slowly and then release their DPA, but cannot grow further.^30,31^ Our TEM observations confirm that the spores lose their DPA (Figure 4C) and Figure S3, and the core appears very discolored, compared to untreated spores Figure (4A). There is also visible degradation of the cortex, spore coat, and exosporium. The level of disruption of the spores varied from still having recognisable spore structural features to very pale outlines with completely unstained internal content. There are major differences from previously reported on spore DPA loss following hypochlorite treatment. In previous studies the spores gradually release their DPA, going from a full DPA signal to no DPA over several minutes. In our case, this process took only 1 minute. It was also not due to temperature or germinants (the spores were suspended in MQ water at 25° C). It is also unlikely that heating from the Raman laser beam catalyzed the reaction since the laser power was only 5 mW, which is comparable to 3 mW used for *B. subtilis* in studies such as Peng et al., 2009.^31^

We attribute the differences observed with the literature where they used similar methods, to different species used in the studies. We used *B. thuringiensis* in our study whereas *B. subtilis* has been used in comparable studies. A major difference is that unlike several other *bacillus* species, *B. subtilis* spores lack an exosporium, which means that its germination mechanics are independent of exosporium damage. This means that sodium hypochlorite acts on the spore coat and cortex, starting its breakdown.^32,33^ In exosporium-producing spores, the exosporium is important in regulating germination,^1,34^ so its degradation can lead to DPA release in *B. thuringiensis*. This effect needs to be thoroughly studied for other *Bacillus* species also. Thus, we conclude from our experiments with 0.5 % sodium hypochlorite that this chemical affects spores Raman spectra by significantly reducing the DPA and DNA peaks within a few minutes up to about 25 minutes. When the DPA release is initiated, it is rapid and within 1 minute all DPA is released. SEM and TEM images together shows significant changes to the exosporium layer, and moderate degradation of the spore coat, cortex, and core. For consistency with other experiments, we verified that spores were inactivated by growing the treated spore suspension and noted a 6 log reduction in CFU.

### Effect of peracetic acid

Peracetic acid is an oxidizing disinfectant agent efficient in inactivating microorganisms via denaturation of proteins, enzymes, and metabolites by oxidation of sulfhydryl and sulfur bonds.^35^ Peracetic acid has been shown to work against spores for some time, and it is effective in solution.^36^ We first measured the Raman spectrum of peracetic acid itself (Figure 1F) and confirmed that it is consistent with previous studies, and that it does not decrease over the measurement time. ^37^ We treated and investigated spores incubated with 1 % peracetic acid. This concentration was chosen as the upper end of the reported sporicidal concentrations of peracetic acid.^38^ As with sodium hypochlorite, there was a variation in the lag time before DPA loss, ranging from 5 minutes to 18 minutes. The speed with which the spores lost the DPA also varied. Of ten spores that were investigated, only two lost their DPA in a minute, similar to the spores treated with sodium hypochlorite, while seven lost the DPA peak in a slower manner, taking from 2 to 10 minutes (Figure 2C). One spore did not lose its DPA over the measurement time. There is a similar downward trend in peak intensity of DNA. This trend continues over the measurement time (30 minutes), which is significantly slower than the reported inactivation time of the spores; at the concentration of 1 % peracetic acid, spores are expected to be inactivated in less than a minute.^10^

When observed using SEM, some spores appeared broken down and degraded, while others were still intact (Figure 3D). In the panel, both a degraded and intact spore in the same field of view is shown. Of the 26 spores imaged and taken from 20 fields of view, 5 were degraded and 21 were intact. It is possible that the degraded spores correlate with the ones that lost their Raman DPA and DNA signal rapidly in the LTRS experiments. This variation observed in the SEM is plausible since spores are heterogeneous.^39^ As with the other experiments, we verified that the treated spores were inactivated by growing the treated spore suspension and noted a 6 log reduction in CFU.

When observed under TEM, spores treated with a peracetic acid showed a clear difference to the untreated spores. In the untreated spores, the spore coat can be seen as several dark layers (Figure 4A), consistent with its dense multilayer structure.^1^ The spore coat in peracetic acid treated spores (Figure 4D and Figure S4) appeared fragmented, separating and breaking into small pieces in all ten imaged spores. The core, cortex, and exosporium appeared intact. This is consistent with the SEM observations as spores with a damaged spore coat can lose their structural integrity. The exosporium and the core did not change visually.

## Conclusion

Rapid detection, whether a spore disinfection procedure was successful or not, are of significance in many areas. We treated *B. thuringiensis* spores with common disinfection chemicals; chlorine dioxide, peracetic acid, and sodium hypochlorite; and measured changes to spores structure and Raman spectra. Chlorine dioxide did not change the Raman spectrum or the spore structure. Peracetic acid shows a time-dependent decrease in the characteristic DNA/DPA peaks and ∼20 % of spores structure were degraded and collapsed, and TEM imaging shows a degradation of the spore coat. Spores treated with sodium hypochlorite show an abrupt drop in DNA and DPA peaks within 20 minutes. The spore structure was overall intact, though internal structural degradation was observed using TEM and the exosporium layer was reduced in size or removed. In all these experiments, structural changes appeared over several minutes, compared to the inactivation time of the spores, which is less than a minute. We conclude that vibrational spectroscopy provides powerful means to detect changes in spores. However, it might be problematic using Raman methods to identify if spores are live or dead directly after a decontamination procedure; no changes in the Raman spectrum occur for chlorine dioxide and changes for the other two chemicals occurs significantly slower than the inactivation process itself.

## Experimental

We used our optical trap and LTRS instrument that is built around a modified inverted microscope (IX71, Olympus).^40,41^ We show an illustration of the system in Figure 5. To trap spores and acquire their corresponding Raman spectra, we use a Gaussian laser beam (TEM00, M2<1.1-1.3) from a continuous wave laser (CRL-DL808-120-S-US-0.5, Crysta-Laser) operating at 808 nm. To prevent laser beam back reflections that might cause laser beam intensity flickering and mode jumping, we use an optical isolator. We pass the laser through a narrow line filter (center wavelength 808 nm, 3.1 nm Bandwidth, Edmund Optic) to improve the Raman signal. To ensure a diffraction-limited spot-size, we spatially filter and expand the beam to fill the back aperture of the microscope objective before coupling the laser into the microscope. We couple the laser into the microscope using a dichroic shortpass mirror with a cut off wavelength of 650 nm. This dichroic mirror serves two purposes, separating the spectrograph side from the conventional imaging side and directing the laser into the microscope objective. We use a 60x water immersion objective (UPlanSApo60xWIR, Olympus) with a numerical aperture of 1.2 and a working distance of 0.28 mm that focuses the beam and forms the trap. With this water immersion objective, we can use the full working distance to position the trap without impacting the trap performance due to spherical aberrations. This freedom in positioning improves signal quality as it allows us to trap objects far away from surfaces that can introduce noise in the form of unwanted Raman scattering and fluorescence.

**Figure 5:**
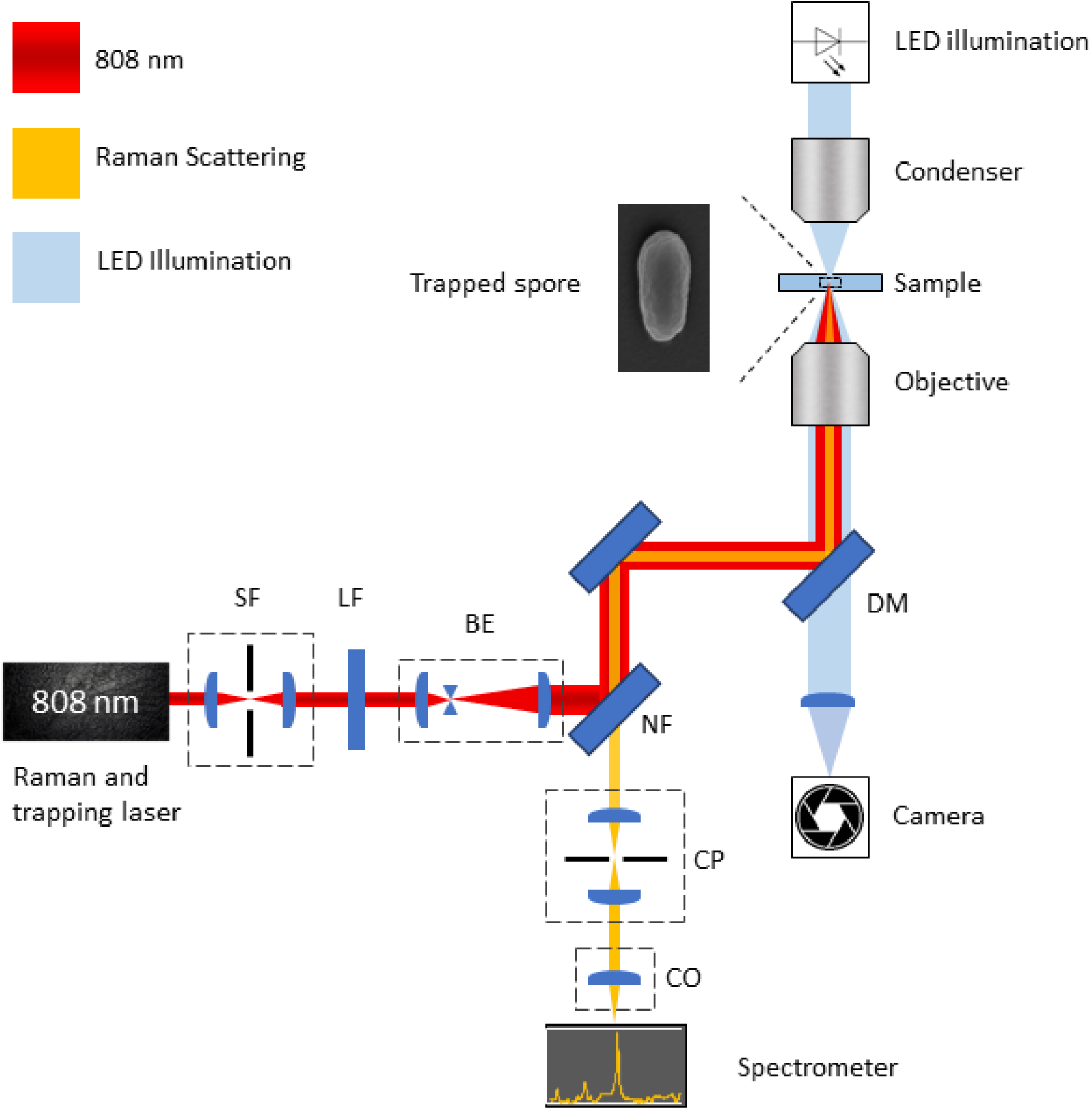
Illustration of the LTRS setup used to aquired Raman spectra of individual spores. The optical system consist of a spatial filter (SF), beam expander (BE), 808 nm line filter (LF), 808 nm notch filter (NF), dichroic 650 shortpass mirror (DM), confocal pinhole (CP), coupling optics for spectrometer (CO). To illuminate a sample we use a LED and acquire images using a CMOS camera.

To perform accurate positioning of the laser focal spot, we aligned the beam so the focal spot coincides with the focal plane of the microscope objective. A diffraction-limited spot of ∼ 800 nm and a depth of field of ∼ 700 nm is created that we can position with nm-resolution in the lateral and axial plane using both a motorized microscope stage (SCAN IM120×100, Märzhauzer Wetzlar) and a piezo stage (PI−P5613CD, Physik Instruments).^42^ Since the objective strongly focuses the laser beam ∼ 65 ° C, the intensity drops fast from the focal spot, both in the lateral and axial directions. We estimate the intensity in the focal spot to 7.81 kW/mm^2^ by measuring the effective power (7 mW) before the objective and using the properties (transmission, focal length, etc.) of the objective from the manufacturer.

To measure the Raman spectrum, we collect the back-scattered light with the microscope objective. First, we pass the back-scattered light through a notch filter (NF808-34, Thorlabs), to remove the Rayleigh scattered light. Then, to maximize the spectral resolution, we expand the beam to fill the spectrometer’s numerical aperture using a telescope. Further, to increase the signal to noise ratio, we place a 150 *μ*m diameter pinhole in the focal point of the telescope, to avoid collecting unwanted light. Finally, we couple the light into our spectrometer (model 207, McPherson) through a 150 *μ*m wide entrance slit where an 800 lp/mm holographic grating disperses the light and the spectrum imaged using a Peltier cooled CCD detector (Newton 920N-BR-DD XW-RECR, Andor) operated at −95 ◦C.

To visualize the sample during measurement, we use an LED lamp with a center wavelength of 470 nm (M470L4, Thorlabs). As the light emitted by the LED has a bandwidth of 26 nm and is located far from the spectral working region of the spectrometer, we can do simultaneous visual imaging and Raman spectroscopy without added spectral noise. To acquire the images, we use a 1920 x 1440 pixel CMOS camera (C11440-10C, Hamamatsu).

To control the sample temperature, we used a home-made software-driven PID regulator, design available upon request. We built the regulator using a micro-controller (STM32F103C8T6, STMicroelectronics), heating foil (Calesco), which we drive using a dual H-bridge (L293D, Texas Instruments), and a thermistor (B57550G1103F005, TDK Electronics) connected through a Wheatstone bridge to a 16-bit ADC (ADS1115, Texas Instruments). To accurately measure the temperature of the sample, we place the thermistor and the heating foil on the nose-cone of the objective, close to the sample. This setup lets us control the sample temperature with sub-mK resolution and an accuracy of 0.1 K. To measure the influence of temperature drifts and noise in general, we use Allan variance to analyze our setup.

### Sample preparation and LTRS measurements

We prepare *B. thuringiensis* (ATCC 35646) spores using BBLTM AK agar 2 Sporulating plates. Spore batches are kept in MQ water at 4°C until use. A stock spore suspension is made with a concentration of 10^6^ spores per ml. To re-suspend the spore stock we vortex them at 2,800 rpm (VM3 Vortex, M. Zipperer GmBH) for 10 sec. A sample is made by adding a 1 cm diameter ring of 1 mm thick vacuum grease on a 24 x 60 mm glass coverslip. Then we add 5 *μl* of the diluted spore suspension inside the ring.

We then proceed to add 5 *μl* of sporicidal chemical on top of the spore suspension, for a final concentration of 750 ppm chlorine dioxide (DKDOX 1500, used at 50 %), 1 % peracetic acid (Acros Organics 35 % stock, diluted 2 % stock prepared on the day of experiment) or 0.5 % sodium hypochlorite (commercial bleach, Klorin brand), respectively, and seal the sample by placing a 23 x 23 mm cover slip on top the grease ring. We immediately start a timer to keep track of how long the spores have been exposed to the chemical. We then place the sample in the LTRS-system and locate a single free floating spore for Raman measurements. The Raman spectrum is recorded using 30 s acquisition time and 2 accumulations per spectra to achieve a temporal resolution of 60 s measured in the range of 600-1400 *cm*^−1^. The spectral resolution is (<2 cm^−1^).

We use a modified version of the above procedure to measure the Raman spectra of spore components (DNA and DPA) and of sporicidal chemicals. Anhydrous DNA and and DPA were sourced from Sigma-Aldrich. Instead of using glass cover slips, a pair 0.25 mm thick quartz cover slips (Alfa Aesar) is used to eliminate the broad Raman peak from the glass. To allow the quartz cover slips to be washed and reused, the grease ring is replaced with a PDMS ring (which can be easily removed). Since there was no particle to trap in these samples, the laser focal point was set 150 *μm* above the quartz surface.

### Verifying viability of treated spores

Treated spore samples were serially diluted and 10 *μl* drops plated onto TSA plates and grown at 30° C overnight. Colonies were counted and compared with the untreated control.

### SEM imaging

To SEM image samples we first prepared a glass cover slip by adding a 20 *μl* drop of 0.1 % poly-l-lysine solution (Sigma-Aldrich) to the cover slip and let the drop evaporate. We marked on the opposite side to make it easier to find the spores. Excessive lysine is removed by gently pouring 2 ml of water to flow over the slide. A 3 *μ*l drop of spore suspension is added on top on the poly-l-lysine drop. When imaging with sporicidal chemicals, we add the chemical on top of the spores and incubate for 30 minutes. Then, the sample is cleaned by again allowing 2 ml of water to flow over the sample to remove the sporicidal chemical and the sample is left to dry completely.

We then coated the sample with a 5 nm layer of platinum, using a Quorum Q150T-ES sputter coater. The samples were then imaged using a Carl Zeiss Merlin FESEM electron microscope to see the spores using InLens and SE-2 imaging modes at a magnification of x50,000.

To ensure that the observed spores are representative of the sample, 20 fields of view were imaged for each sample.

### TEM imaging

Samples for TEM were prepared as liquid suspensions of spores after 30 minutes treatment with peracetic acid, sodium hypochlorite and chlorine dioxide as before, as well as an untreated sample suspended in water. After the incubation, samples with sporicidal chemicals were centrifuged and resuspended in MQ water to wash off the chemical.

Spores were fixed with 2.5 % Glutaraldehyde (TAAB Laboratories, Aldermaston, England) in 0.1M PHEM buffer and further postfixed in 1 % aqueous osmium tetroxide. They were further dehydrated in ethanol, acetone and finally embedded in Spurr’s resin (TAAB Laboratories, Aldermaston, England). 70nm ultrathin sections were post contrasted in uranyl acetate and Reynolds lead citrate. Samples were examined using a Talos L120C (FEI, Eindhoven, The Netherlands) operating at 120kV. Micrographs were acquired with a Ceta 16M CCD camera (FEI, Eindhoven, The Netherlands) using TEM Image and Analysis software ver. 4.17 (FEI, Eindhoven, The Netherlands).

## Acknowledgement

This work was supported by the Swedish Research Council (2019-04016) and from the Kempestiftelserna (JCK-1916.2). We thank Anna-Lena Johansson at FOI for preparing spores in this project.

The authors acknowledge the facilities and technical assistance of the Umeå Core Facility for Electron Microscopy (UCEM) at the Chemical Biological Centre (KBC), Umeå University, a part of the National Microscopy Infrastructure NMI (VR-RFI 2016-00968)

