## Supplemental information for "Disinfection chemicals mode of action on the bacterial spore structure and their Raman spectra"

**Correspondence**

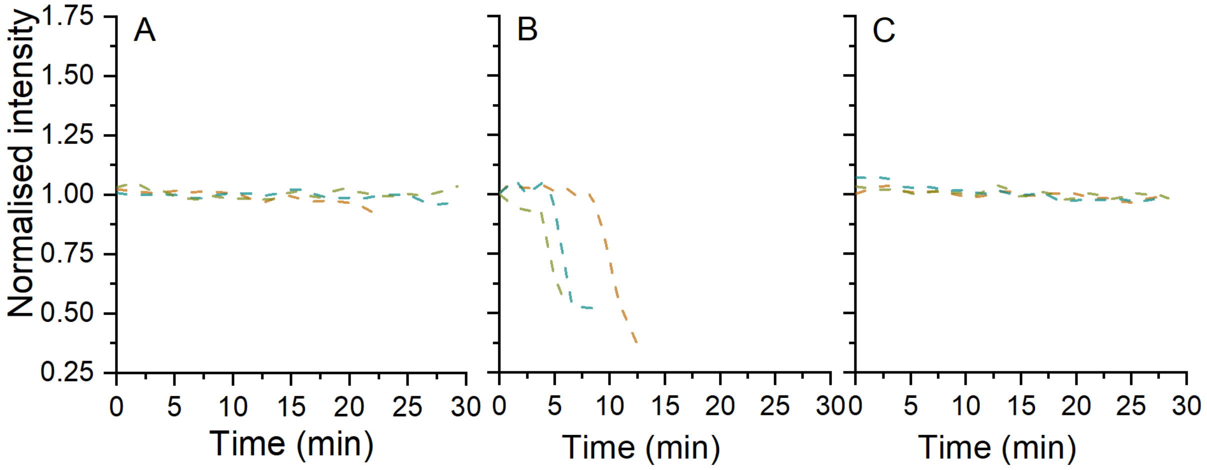


Figure S1. Changes in the normalised intensity of the 1580 $cm^{-1}$ (Amide) peak over time with chlorine dioxide (A), sodium hypochlorite (B) and peracetic acid (C) (n=3 for each chemical). Aside from the Raman peak studied, the experimental conditions are the same as those used in Figure 2. There was no reduction in the peak for chlorine dioxide or peracetic acid.

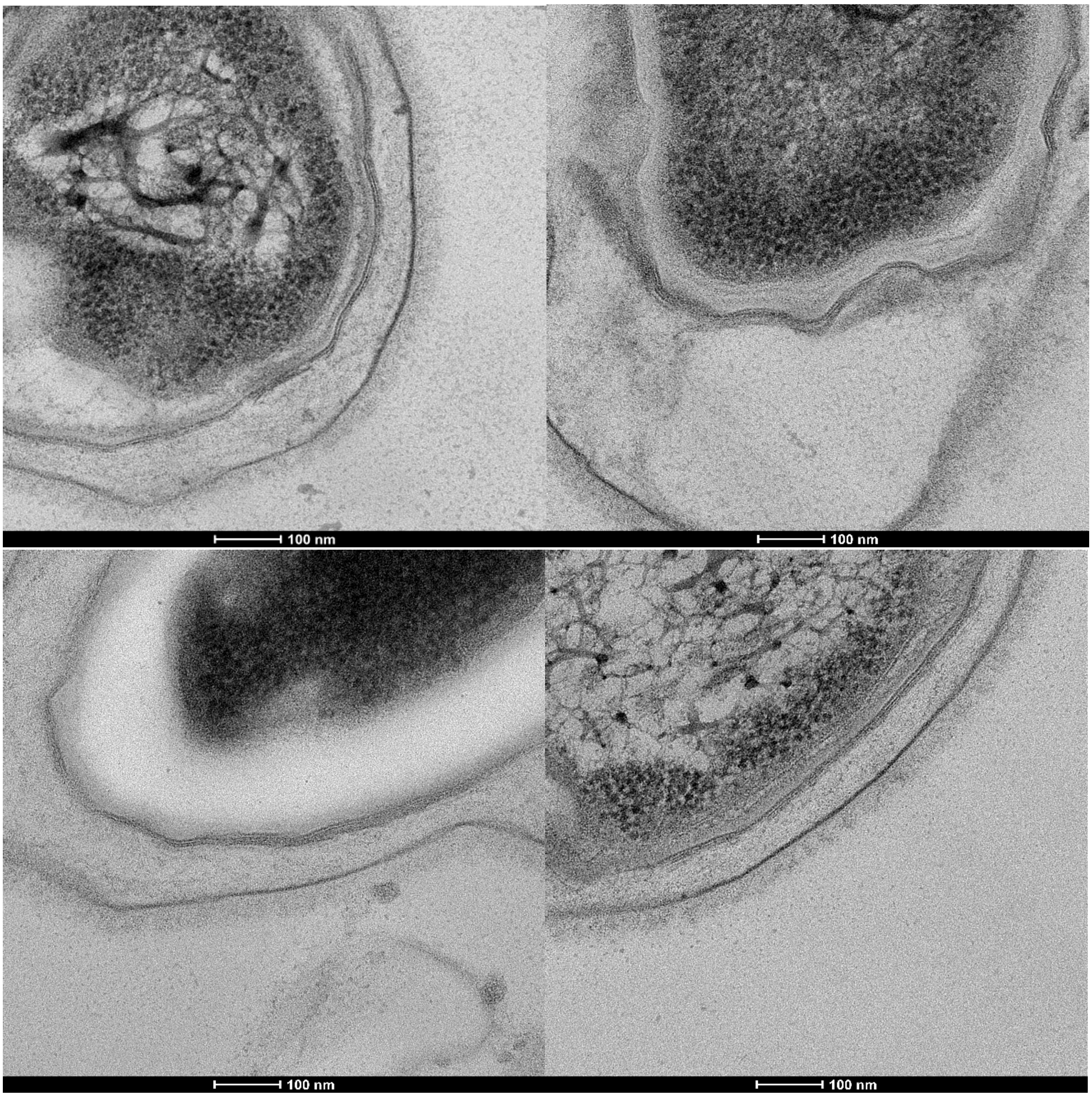


Figure S2. Additional fields of view of untreated *B. thuringiensis* spores. Note the continuous unbroken lines of the spore coat layers.

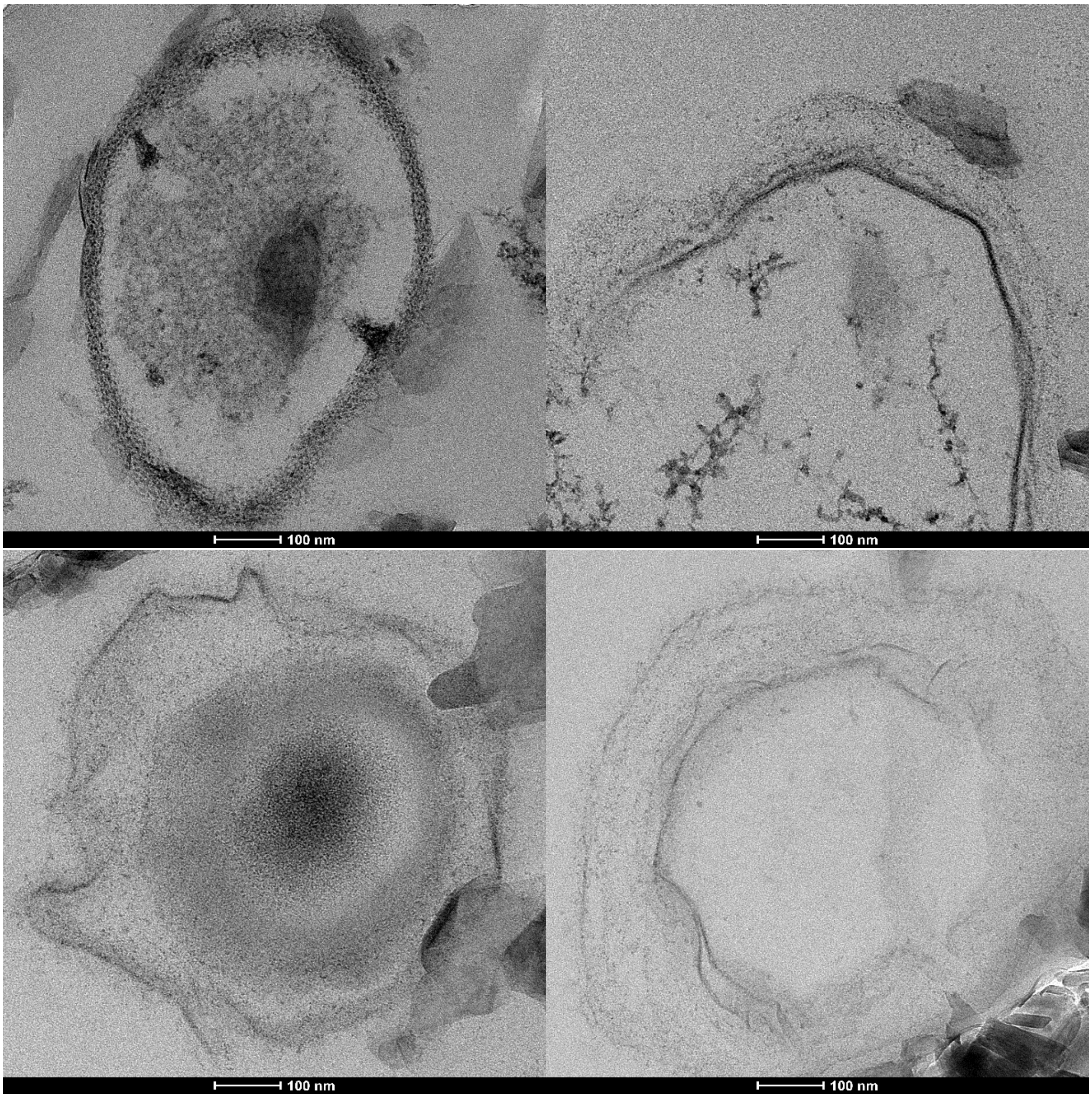


Figure S3. Additional fields of view showing spores degraded by sodium hypochlorite.

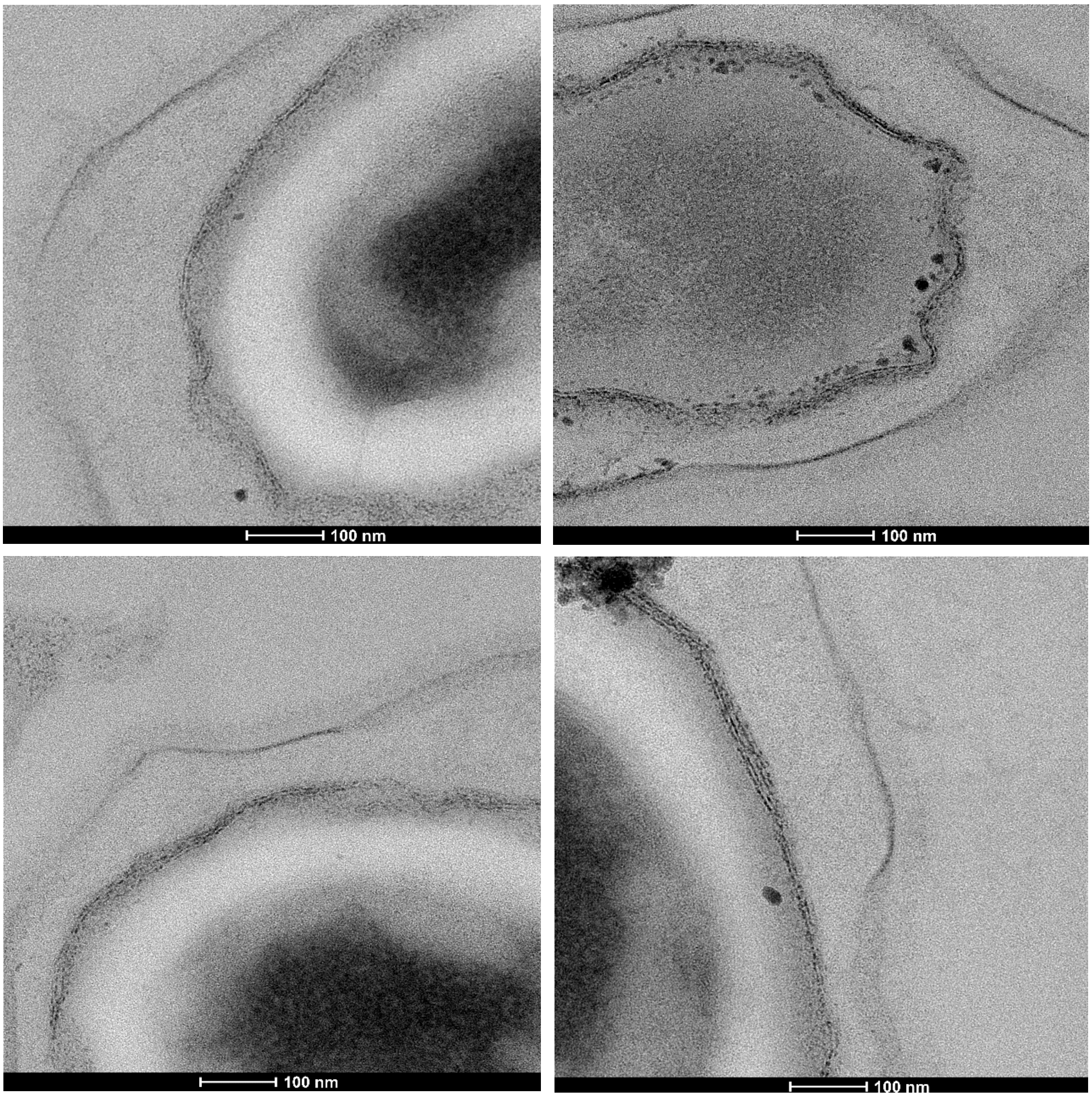


Figure S4. Additional fields of view showing damage to the spore coat from incubation with 1% peracetic acid. Note the fragmentation of the spore coat.
